# Ovarian embryonal rhabdomyosarcoma is a rare manifestation of the DICER1 syndrome

**DOI:** 10.1101/011304

**Authors:** Leanne de Kock, Harriet Druker, Evan Weber, Nancy Hamel, Jeffery Traubici, David Malkin, Jocelyne Arseneau, Dorothée Bouron-Dal Soglio, John R. Priest, William D. Foulkes

## Abstract

Embryonal rhabdomyosarcoma (ERMS), a malignant soft tissue sarcoma, is one of the most common paediatric cancers. Certain ERMS tumours are associated with the DICER1 syndrome, a distinctive tumour predisposition syndrome caused by germ-line mutations in the microRNA-maturation pathway gene, *DICER1*. In addition to germ-line *DICER1* mutations, highly characteristic somatic mutations have been identified in several DICER1-associated tumour types. These so-called “hotspot” mutations affect highly conserved amino acid residues central to the catalytic activity of the DICER1 ribonuclease IIIb domain.

Primary ovarian ERMS (oERMS) is extremely rare. We present a case of a 6-year-old girl with an oERMS found to harbour two mutations in *DICER1*. In addition to the oERMS, the girl also exhibited other DICER1 phenotypes, including cystic nephroma (CN) and multinodular goitre. Somatic investigations of the CN revealed the presence of a hotspot *DICER1* mutation different from that in the oERMS. Of particular interest is the CN presented at the age of 12 years, which is much older than previously reported age range of susceptibility (birth to four years of age).

This report documents both germ-line and highly characteristic somatic *DICER1* mutations in a case of oERMS, adding to the expanding spectrum of rare childhood tumours in the DICER1 syndrome.

## Introduction

Embryonal rhabdomyosarcoma (ERMS) is a malignant tumour of mesenchymal origin and one of the most common extra-cranial solid tumours of childhood. ERMS tends to affect children within the first decade of life and the more common sites are orbit, nasopharynx, vagina and bladder. The majority of ERMS arise sporadically, but associations between ERMS and inherited familial syndromes, such as the Li-Fraumeni syndrome, neurofibromatosis and Beckwith-Weidemann syndrome suggest that a genetic predisposition contributes to some cases^1–5^.

Recently it has become evident that some ERMS are associated with germ-line *DICER1* mutations. In general, *DICER1* mutations cause a distinctive tumour predisposition syndrome, the DICER1 syndrome, (OMIM # 601200) which is characterized by pleuropulmonary blastoma (PPB) and other rare childhood neoplasms^6^. PPB is a mixed pattern sarcoma in which ERMS is a predominant histologic subtype^7^. *DICER1* mutations are strongly associated with ERMS of the uterine cervix (cERMS), a generally uncommon site for ERMS, which occurs from the age of approximately 10-20 years^5, 8, 9^. Both germ-line and highly characteristic somatic “hotspot” *DICER1* mutations have been reported in two cERMS^8, 10^ and bladder ERMS has been observed in two *DICER1* kindred^7, 11^. *DICER1* mutations were identified in two of 52 sporadic ERMS cases^7^. No associations are evident between *DICER1* mutations and alveolar rhabdomyosarcoma (ARMS), a rhabdomyosarcoma subtype tending to affect older children and believed to arise through biological mechanisms distinct from ERMS^12^.

Like cERMS, ovarian ERMS (oERMS) is particularly rare^13^; a case of oERMS, which was histologically similar to cERMS and potentially associated with a *DICER1* mutation, has been mentioned in the literature^5, 14^, but the child’s mutation status was not reported. We report detailed investigation of *DICER1* gene in a six-year-old girl with oERMS and other *DICER1* phenotypes.

## Materials and Methods

The study was approved by the Institutional Review Board of the Faculty of Medicine of McGill University no. A12-M117-11A and was performed with full informed parental consent. The tumours were reviewed by pathologists at the institution from which the samples were acquired and by central reference pathologists (JA and DB-DS).

### Molecular Screening of DICER1

Somatic “hotspot” mutations affecting the RNase IIIa and RNase IIIb domains were screened for by PCR amplification of gDNA extracted from each of the formalin-fixed, paraffin-embedded (FFPE) tumours, followed by Sanger sequencing (McGill University and Genome Quebec Innovation Centre (MUGQIC)), as previously described^15–17^.

### Immunohistochemistry (IHC)

Immunostaining of desmin, myogenin and MyoD1 was performed on deparaffinised 4 μm tissue sections using a DAKO Auto-stainer Link 48, following the manufacturer’s instructions with only minor modifications (Dako Autostainer 48, CA). The MyoD1mouse monoclonal antibody (Dako) was used at a dilution of 1:50. The desmin mouse monoclonal antibody (Dako) and Myogenin mouse monoclonal antibody (Dako) were used without further dilution.

## Case Report and results

A six-year-old girl of Ashkenazi Jewish and Anglo-Saxon descent presented with a three month history of abdominal distension, rigidity and difficulty passing urine. A mass on the right ovary was discovered, measuring 11.5 cm x 10 cm x 7 cm (Fig. 1a). Surgical resection of the right ovarian tumour, right fallopian tube and part of the omentum was performed and she was treated with antinomycin D, vincristine and cyclophosphamide in conjunction with etoposide and ifosfamide therapy. She also received whole abdomen radiation 1800 cGy in 12 fractions. Pathology of the multiloculated mass showed variability in architecture and differentiation: an edematous myxoid matrix containing small pleomorphic spindle cells constituted the poorly differentiated areas of the tumour. The tumour cells showed evidence of skeletal muscle differentiation. The more well-differentiated areas of the tumour contained larger cells with abundant eosinophilic cytoplasm and oval to indented nuclei. Islands of cartilaginous differentiation were focally present within the tumour and several mitotic figures were noticeable (Fig. 2a, panel I and II). The capsule of the ovary was also focally thickened by a fibroblastic proliferation. Five years post-radiation, she developed radiological focal nodular liver hyperplasia. At 12 years of age, she developed a cystic nephroma (CN) of the left kidney (Fig. 1b, panel II) and underwent a partial nephrectomy. At 13 years of age, she developed multinodular goitre (MNG) (Supplementary Figure S1 and S2) and at 16 years of age she developed a fibroadenoma of the breast.

**Figure 1 Legend:**
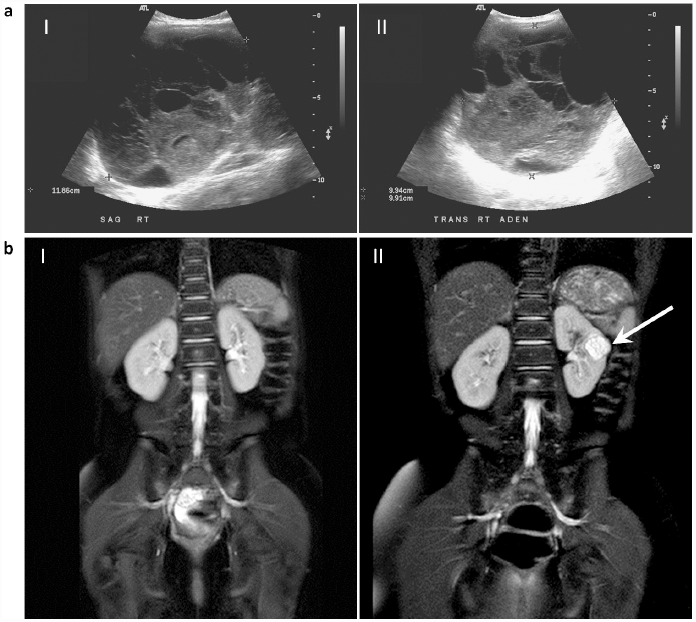
**a)** Sagittal **(panel I)** and transverse **(panel II)** ultrasound images of right ovary showing multiloculated ovarian tumour; taken at 6-years, 3-months of age. **b)** Magnetic resonance image (MRI) taken at 11.5 years of age revealed no renal abnormalities **(panel I)**. Seven months later, at 12 years of age, a 21mm cystic mass (indicated by arrow) was identified on the left kidney on MRI **(panel II)**.

**Figure 2 Legend:**
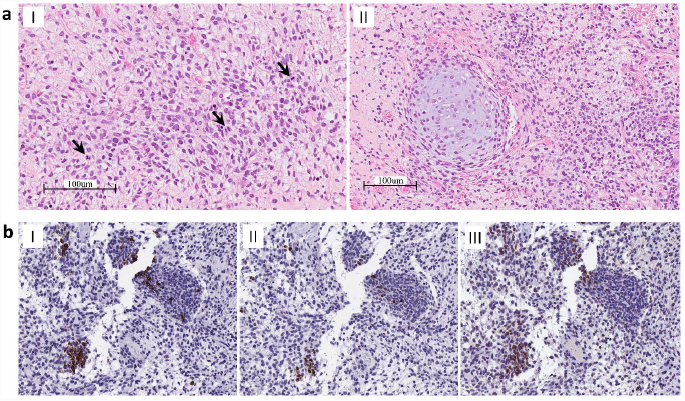
**a) Panel I:** Small pleomorphic spindle cells in a slightly myxoid matrix. Notice several mitotic figures (indicated by arrows). Panel II: Island of cartilaginous differentiation. **b)** Immunohistochemical staining of desmin **(panel I)**, myogenin **(panel II)** and MyoD1 **(panel III)** showed focal positivity within the tumour cells for each protein (200 x magnification).

The rarity of oERMS tumours makes their definitive diagnosis challenging. A full diagnostic workup of the small, round blue cell tumour was thus performed using a number of ancillary tests. Light and electron microscopy revealed skeletal muscle differentiation. Immunohistochemical (IHC) staining of myogenesis-associated proteins including desmin, myogenin and MyoD1 was performed – focal positivity for all three proteins was evident in the tumour cells (Fig. 2b, Panels I to III). Molecular genetic analyses were performed at the primary institution to test for the presence of the t(1;13) and t(2;13) reciprocal translocations known to be associated with ARMS^4^ - No hybrid transcripts were detected for either translocation. Alternate diagnoses were considered during the differential diagnosis including a teratoma or poorly differentiated Sertoli-Leydig cell tumour with heterologous elements. However, no endodermal or ectodermal elements were found to support the former diagnosis, and no convincing Sertoli cell or Leydig cell differentiation was identified. Following thorough pathological review, ERMS of the ovary was the final diagnosis.

Molecular genetic sequencing of the patient’s germ-line gDNA revealed a deleterious mutation, c.1196_1197dupAG, in exon 8 of *DICER1* (Prevention Genetics, Marshfield, WI, USA). This mutation is predicted to cause a translational frame-shift which would induce a premature stop codon within the sequence coding for the trans-activating response RNA-binding protein binding domain (TRBP-BD) [p.(Trp400Serfs*59)]. Translation of this mutant transcript, if not degraded by nonsense-mediated decay (NMD), would result in the expression of a severely truncated protein consisting of only the DExD/H box helicase domain and a portion of the TRBP-BD (Fig. 3a). We confirmed this mutation and also identified an acquired somatic mutation in the oERMS tumour gDNA: c.5425G>A mutation, which is predicted to result in p.(Gly1809Arg) at the protein level (Fig. 3b, Panel I). A different somatic *DICER1* mutation, c.5439G>T, predicted to cause p.(Glu1813Asp) at the protein level, was identified in the CN tumour gDNA (Fig. 3b, Panel III). Both somatic mutations are predicted to be damaging by both SIFT and PolyPhen2 with respective scores of 0 [0.02 for the p.(Gly1809Arg) mutation] and 1. No tissue was available for molecular analysis from the MNG or the fibroadenoma.

**Figure 3 Legend:**
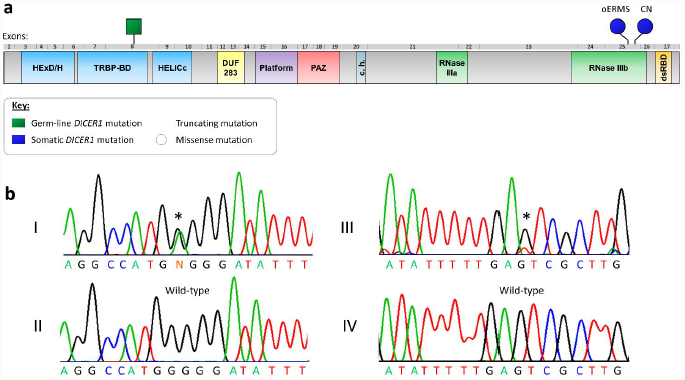
**a)** Graphic representation of the unfolded DICER1 protein, indicating the approximate positions of the p.(Trp400Serfs*59) (germ-line), p.(Gly1809Arg) (somatic – oERMS) and p.(Glu1813Asp) (somatic – CN) amino acid changes. DICER1 domains defined as follows: DExD/H, DExD/H box helicase domain; TRBP-BD, trans-activating response RNA-binding protein binding domain; HELICc, helicase conserved C-terminal domain; DUF283, domain of unknown function; Platform, platform domain; PAZ, polyubiquitin associated zinc finger domain; c.h., connector helix; RNase IIIa, Ribonuclease IIIa domain; RNase IIIb, Ribonuclease IIIb domain; dsRBD, double-stranded RNA-binding domain. **b)** Chromatogram showing the somatic c.5425G>A mutation in gDNA extracted from the oERMS **(panel I)**. The wild-type sequence is illustrated in **panel II**. The somatic c.5439G>T mutation identified in gDNA extracted from the CN is presented in **panel III** compared to the wild-type sequence in **panel IV**.

## Discussion

Ovarian tumours associated with the *DICER1* syndrome are predominantly non-epithelial in origin. SLCT (a non-epithelial ovarian sex-cord stromal tumour) is the most frequently observed ovarian tumour in *DICER1* mutation carriers^14, 15^ and somatic *DICER1* missense mutations occurring within the RNase IIIb domain have been identified in 60% of ovarian SLCTs^10, 15^. Other ovarian sex cord-stromal tumours reported with a *DICER1* mutation include juvenile granulosa cell tumour and gynandroblastoma^14, 15^. The age of onset for *DICER1*-associated SLCT typically ranges from 2 to 45 years of age with a peak onset between 10 and 15 years of age^6^. We sought to determine the approximate range for age of susceptibility for oERMS tumours. Our literature search (summarised in Supplementary Table S1) revealed 32 well-documented reports of ovarian rhabdomyosarcomas, nineteen of which are of the embryonal subtype. The ages at first symptom report of the ERMS cases ranged from 6 to 86 years of age, with a median age of 25 years. The oERMS case we report presented at 6 years of age. It remains to be seen whether oERMS occurring in association with a *DICER1* mutation tend to present at an earlier age, as is the case for the majority of the DICER1-associated diseases^6^. The cystic nephroma, a common DICER1 phenoptype, in the patient we report is particularly notable because the 2.1 cm-diameter CN is documented to have developed at age 12 years during a 7-month interval between surveillance magnetic resonance scans (Fig. 1b, panels I and II). Almost all DICER1-associated CN are diagnosed before age 4 years^18, 19^; in rare CN cases diagnosed at an older age, the age at which the CN actually developed is uncertain. Documented development of CN at age 12 years is unprecedented and illustrates the challenges of disease screening in *DICER1* mutation carriers. The CN contained a somatic RNase IIIb mutation, discussed below, which is typical for CN^20^. This child had received 1800 cGy total abdominal radiation 6 years prior to development of CN. There is, however, no suggestion in the literature that therapeutic doses of ionizing radiation might predispose to development of CN or to DICER1 RNase IIIb mutations.

The somatic mutations identified within the oERMS (c.5439G>T) and CN (c.5425G>A) tumour samples are typical of somatic mutations in other *DICER1* phenotypes^6^ and occurred within the sequence encoding the RNase IIIb domain of DICER1, affecting highly conserved amino acid residues. The DICER1 protein is a small RNA processing endoribonuclease, which measures and cleaves microRNA (miRNA) hairpin precursors to generate mature miRNAs, which subsequently regulate the translation of target-gene mRNA transcripts. The amino acids altered by the mutations (Gly1809 and Glu1813) are metal-ion binding residues central to the cleavage of 5p miRNAs from the 5′ arm of miRNA precursors (pre-miRNAs). Mutations of these RNase IIIb sites disrupt the processing of pre-miRNAs and consequently reduce the population of 5p miRNAs within the cell^21, 22^. As has recently been reported^23^, we noted the occurrence of two different somatic *DICER1* RNase IIIb mutations in two tumours that arose in the patient.

oERMS has striking histologic similarities, including primitive cartilage elements, to other distinctive DICER1-related tumours like PPB and cERMS^5^, which points to a common histological origin and genetic cause. The discovery of a somatic RNase IIIb *DICER1* mutation in an oERMS arising in a germ-line mutation carrier adds to the growing literature that these RNase IIIb mutations are key mutational events in DICER1-related tumourigenesis. The downstream perturbations of miRNA biogenesis that arise in cells in which DICER1 function is impaired are currently being explored and the underlying mechanisms of tumourigenesis are being investigated. Sequencing of *DICER1* in additional oERMS tumours may further substantiate this tumours’ association with the DICER1 syndrome.

## Conclusion

This report documents both germ-line and highly characteristic somatic *DICER1* mutations in a case of oERMS, suggesting that oERMS is among the growing list of rare childhood tumours associated with the DICER1 syndrome. Screening for *DICER1* mutations should be considered for children presenting with this rare condition. This report also documents an unusually late presentation of CN at the age of 12 years. Taken together with the primary finding of an oERMS, this finding shows that the clinical phenotype associated with *DICER1* mutations remains unsettled.

## Acknowledgements

We thank the family involved in this research for their consent to participation and the clinicians for referring the case and providing samples. We also thank Dr. Steffen Albrecht for assistance with the figures. This research was made possible thanks to the support of the Alex’s Lemonade Stand Foundation grant to Dr. William D. Foulkes, and the CIHR/FRSQ training grant in cancer research FRN53888 of the McGill Integrated Cancer Research Training Program and the McGill Faculty of Medicine Internal Studentship Award (Awarded as a Ruth and Alex Dworkin Fellowship and a Graduate Excellence Fellowship) to Leanne de Kock.

